# Agricultural impacts on streams near Nitrate Vulnerable Zones: a case study in the Ebro basin, northern Spain

**DOI:** 10.1101/663054

**Authors:** Rubén Ladrera, Oscar Belmar, Rafael Tomás, Narcís Prat, Miguel Cañedo-Argüelles

## Abstract

Agricultural intensification during the last century has produced river degradation across Europe. From the wide range of pressures derived from agricultural activities that impact rivers, diffuse agricultural pollution has received most of the attention from managers and scientists. The aim of this study was to determine the main pressures exerted by intensive agriculture around Nitrate Vulnerable Zones (NVZs), which are areas of land that drain into waters polluted by nitrates according to the European Nitrate Directive (91/676/EEC). The study area was located in the NW of La Rioja (Northern Spain), which has the highest levels of nitrate concentrations within the Ebro basin. The relationships between forty environmental variables and the taxonomic and functional characteristics of macroinvertebrate assemblages (which are good indicators of water quality) were analyzed in 11 stream reaches differentially affected by upstream agricultural activity. The streams affected by a high percentage of agricultural area had significantly greater nitrate concentrations and distinct macroinvertebrate assemblages dominated by pollution tolerant taxa. Hydromorphological alteration (i.e. channel simplification, riparian forest degradation and sediment inputs), which is closely linked to agricultural practices, was the main factor affecting macroinvertebrate assemblages. Good agricultural practices should be implemented in streams affected by NVZs to reverse stream degradation, in consonance with the European Water Framework Directive (WFD). Management actions in these areas should not focus exclusively on nitrate reduction, but also on restoring riparian and aquatic habitats.

## Introduction

Agriculture is the most widespread form of land-use change [1,2], and it has been identified as the main driver of global biodiversity decline [3] and one of the most important stressors affecting freshwater ecosystems in Europe [4] and around the world [2]. Water quality is strongly affected by agrochemicals, pesticides and nutrients intensively used in productive agriculture [5]. Among them, nitrate is one of the most abundant agrochemical pollutants in waters of rural areas [6], since it is intensively used for plant growth and development [7]. Intensive agriculture not only needs agrochemicals but also important water quantities to increase productivity. This need for water has led to strong river and stream regulation in the last century that has reduced and altered (e.g., inversed) flow [8–10]. Finally, agriculture degrades the fluvial habitat through riparian forest removal, channel incision and rectification, reduction of bank stability and sediment deposition [11].

There is a broad bibliography on the impact of agricultural activity on rivers based on macroinvertebrate assemblage [e.g. 11–17], with different studies specifically assessing the impact of nitrate on macroinvertebrates [see 18,19]. Nitrate impact in rivers communities has been especially associated with increased autotrophic production favored by increases in light availability due to riparian vegetation loss [20,21]. High periphyton biomass reduces diurnal oxygen concentrations, impacting on pollution sensitive species such as some Ephemeroptera, Plecoptera and Trichoptera (EPT) and reducing substrate availability for grazing mayflies, which are replaced by molluscs, oligochaetes and chironomids [22].

Beyond this nitrate toxicity and algal growth, stream macroinvertebrate assemblages located in agricultural areas are especially sensitive to habitat deterioration, which is mainly related with an increase in fine sediments [23–25]. Fine sediments remove and homogenize habitats [25], reduce taxa richness, especially some EPT species [24,25], and alter assemblage composition [26]. They also alter macroinvertebrate functional composition, which has proved to be an interesting tool to determine agricultural impacts in rivers [15,17,23,27–29]. Higher amounts of sediment can cover vegetal and mineral coarse substrate leading to an increase in burrowers and grazers, and a decrease in crawlers and scrapers [15,17,28,29].

Different Directives have been implemented in the European Union (EU) in the last decades to revert agricultural impacts on rivers, as the Nitrates Directive (91/676/EEC; ND), which was adopted in 1991. The ND aims to protect water quality across Europe reducing nitrates from agricultural sources and promoting the use of good farming practices. Within the ND, Nitrate Vulnerable Zones (NVZs) are defined as the areas of land that drain into waters polluted by nitrates, and farmers with land in these zones have to follow mandatory rules to recover water quality [30]. The ND forms an integral part of the European Water Framework Directive (2000/60/EC; WFD), which is aimed at establishing a framework for water protection, so all the water bodies in Europe reach a “good ecological status” by 2021 or 2027. To achieve this good ecological status, the WFD indicates that pressures and impacts at basin scale must be identified to guide management measures. This identification is often inadequate in existing monitoring programs [31].

This study assesses the potential impact of agriculture on stream ecosystems around NVZs where nitrate pollution has been pointed out as the main existing pressure using the taxonomic and functional composition of macroinvertebrate assemblages as indicators. The Ebro basin was selected as a case study because of being subjected to heavy human pressures, especially agriculture [32,33]. The specific objectives of the study were: i) determining the main agricultural pressures on rivers located in areas affected by intensive agriculture around NVZs; and ii) assessing their degree of compliance with WFD according to physicochemical variables and macroinvertebrate indices. We hypothesized that these streams would be affected not only by nitrate pollution, but also by an increase in fine sediments and a reduction in riparian and aquatic habitat quality. As a consequence, we expected a shift in the functional structure and taxonomic composition of aquatic macroinvertebrate assemblages, from a domination of pollution sensitive species feeding mainly on coarse organic matter (e.g. vegetal detritus) to a domination of tolerant species feeding mainly on fine organic matter (e.g. suspended particles) and algae.

## Methodology

### Study area and sampling strategy

The study was carried out in 6 first-order streams located in the NW Spain (Fig 1), in the Oja-Tirón, Zamaca and Najerilla sub-basins (belonging to the Ebro basin). The 6 streams belong to the typology “Calcareous mediterranean mountain rivers” according to the classification developed in Spain to fulfil the WFD (ORDER ARM/2656/2008, Ministry for the Environment). Eleven sites without important urban or industrial affections were studied, mainly differing in the upstream agrarian activity (i.e. percentage of cultivated land). Stream sections affected by other anthropogenic pressures were not included to avoid potentially interfering factors.

**Fig 1.**
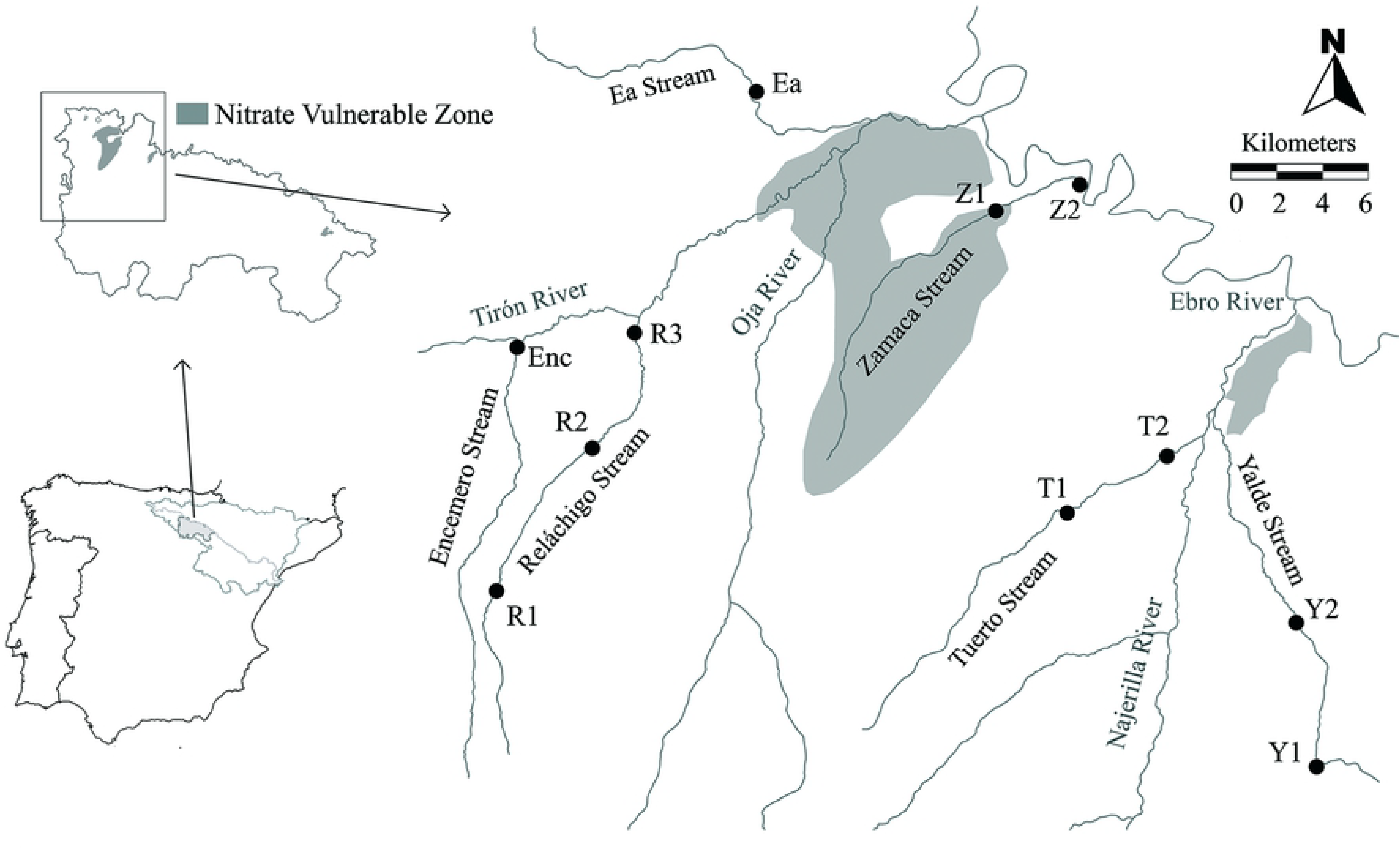
Map of the study area. Sampling sites in the study area (La Rioja, Spain). Nitrate Vulnerable Zones (NVZs) are coloured in grey (“Zamaca and Oja alluvial” and “Low Najerilla alluvial”).

Well-conserved forests exist in the upper course of Yalde and Reláchigo Streams (Fig 1), where agricultural activity is scarce. The cultivated area increases as the streams flow into the Ebro Valley, where intensive farming dominates (mainly cereal, vineyard and horticultural crops). As a result of the intense agricultural activity, two NVZs exist in the study area, called “Zamaca and Oja alluvial” and “Low Najerilla alluvial”.

Each site was sampled 4 times during 2017 (i.e. February, May, August and November). Physicochemical variables were analyzed at each sampling occasion, whereas habitat quality and macroinvertebrates were studied exclusively in May.

### Environmental variables

Forty environmental variables belonging to different categories (physicochemical, hydromorphological, geological, land cover and topographic; S1 Table) were studied at each site.

Water temperature (°C), pH, dissolved oxygen (ppm) and electrical conductivity (mS/cm) were measured *in situ* using selective electrodes. For ion analysis, 125 ml water samples were collected and transported in a cool-box to the laboratory, where they were frozen until being analyzed using chromatographic standard methods [34].

The riparian habitat was characterized using the four components of the QBR Riparian Forest Quality Index by Munné *et al.* [35]: total riparian vegetation cover, cover structure, cover quality and channel alterations. The fluvial instream habitat was characterized using the IHF River Habitat Index [36], which measures habitat heterogeneity by analyzing seven features: substrate embeddedness, rapid frequency, substrate composition, velocity/depth conditions, percentage of shading, heterogeneity components and in-channel vegetation cover. Additionally, percentage of macrophyte cover was visually determined.

The geological, land cover and topographic variables were derived, respectively, though layers from the Spanish Geological Survey (IGME), the Spanish Land Cover and Use Information System (SIOSE) and the Spanish National Geographic Institute (IGN) using a Geographic Information System (GIS).

### Macroinvertebrate sampling and biotic indices

Macroinvertebrates were collected according to the Standard protocol for benthic invertebrates of wadeable streams in Spain (ML-Rv-I-2013; [37]). The material was preserved in 70 % ethanol and taken to the laboratory to be identified. The identification of macroinvertebrates was generally done to genus level, except for some Diptera families, Gammaridae, Oligochaeta and Hydracharina. If necessary, sub-sampling was conducted to estimate the taxa abundances, and at least 300 individuals per sample were identified and counted.

IBMWP [38] and IMMi-T [39] biotic indices were calculated using the MAQBIR software [40]. The IBMWP (Iberian Biological Monitoring Working Party) is based on the tolerance of macroinvertebrate families to water quality and river alteration. The presence of each family provides a single score out of 10 (being 1 highly tolerant and 10 highly sensitive) with the cumulative scores providing the final IBMWP score. The IMMi-T (Iberian Mediterranean Multimetric Index, using quantitative data) is a multimetric index based on the number of families, EPT, IASPT and log (selected EPTCD + 1). Following the WFD, this index was standardized by calculating the EQR value (Ecological Quality Ratio) dividing the value of each metric by the reference value for Calcareous Mediterranean mountain rivers.

### Biological traits

Seven biological traits (microhabitat, locomotion, trophic status, reproductive cycles, respiration, food and feeding habits) containing 45 categories (S2 Table) obtained from a database [41] were used to describe macroinvertebrate functional structure. The trait categories in this database have an affinity score assigned for each taxa ranging from 0 to 5, from null to high affinity, respectively [42]. To analyze the functional structure, a dataset of relative abundance of trait categories per sample was built by using a ‘fuzzy coding’ approach [42].

### Statistical analysis

In order to determine the main environmental variables related to differences in macroinvertebrate assemblage among sites, a DISTLM analysis was performed (PERMANOVA + for PRIMER, [43]) based on genus abundance. The macroinvertebrate distance matrix was created using the Bray Curtis distance method after the assemblages’ data were log (*x* + 1) transformed. The environmental variables were log (*x* + 1) transformed and normalized. The values of physicoquemical variables included in the analysis were the averages of the four sampling dates since they represent better the physicochemical state of the streams and their influence in the biological assemblages. The DISTLM routine was done using a step-wise selection procedure based on the AIC selection criteria [44], to obtain a set of variables capable to explain changes in macroinvertebrate composition.

Two groups of sampling sites were differentiated based on a hierarchical cluster analysis of the macroinvertebrate assemblages (unweighted pair group method using arithmetic averages, UPGMA; PRIMER, [45]). Then, an Indicator Species Analysis (IndVal, [46]) was used to determine the taxa significantly associated with each of the groups. This method provides an indicator value (IV-value) for each taxon according to its presence and abundance in each group. A Monte Carlo permutation test with 9999 permutations was used to test the significance of each IV-value (P<0.05). For each biological trait category, a nonparametric pairwise Mann-Whitney U test was performed to detect differences between the groups defined in the cluster analysis. Asymmetric beanplots (software R3.4.2, package beanplot, [47]) were used to visualize differences in those biological traits that resulted significantly different between groups.

## Results

### Environmental variables, hydromorphological and biotic indices and WFD compliance

The average oxygen concentrations of the different studied sites varied between 6.90 and 10.17 mg/L, registered at Y2 and R1 respectively (Table 1). Every site and date showed oxygen concentrations greater than 5 mg/L (the lowest value established by RD 817/2015 - Spanish legislation based on WFD - to consider a river in good status), except T2 on august, where oxygen concentration was 4.30 mg/L.

**Table 1.**
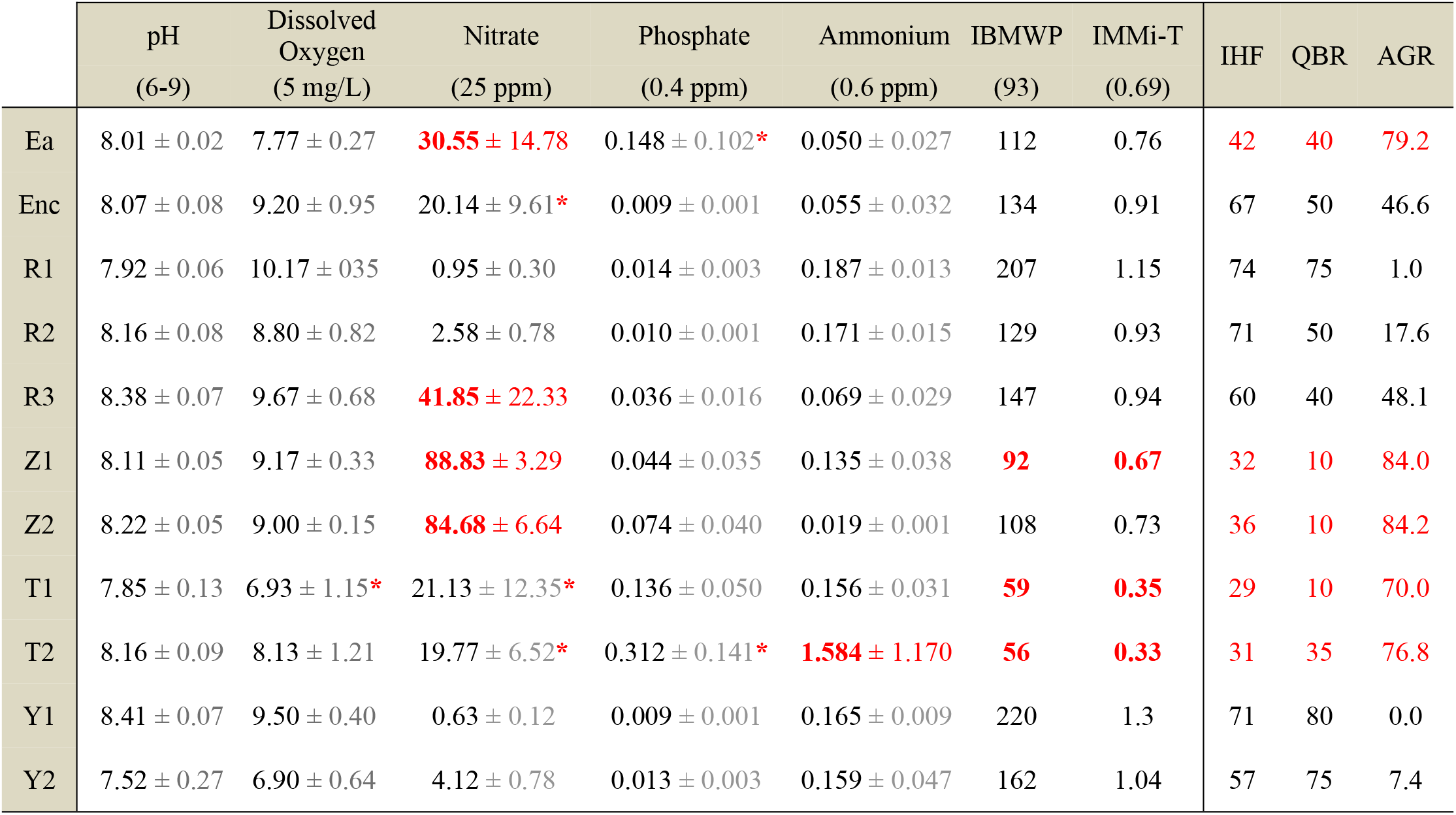
Environmental variables. Values of the physicochemical variables and the biotic indices -IBMWP and IMMi-T-used to assess the good ecological status of streams according to Spanish legislation (RD 817/2015), based on the European Water Framework Directive (WFD). The limits established by Spanish legislation to consider a water body at good ecological status or higher for “Calcareous Mediterranean mountain rivers” are shown in parentheses. For chemical variables, four sampling average values ± SE are shown. Values not achieving good ecological status according to RD 817/2015 are highlighted in red bold. Red asterisk indicate that this parameter did not achieve the good status every sampling date, although average value achieved. IHF and QBR index values, which are not considered by RD 817/2015 to determine the good ecological status of a water body, are shown at the right part along with the percentage of area occupied by agricultural land upstream the river reach (AGR). The lowest values of QBR (<50) and IHF (<50) and the highest values of AGR (≥70) are highlighted in red.

|  | pH<br>(6-9) | Dissolved<br>Oxygen<br>(5 mg/L) | Nitrate<br>(25 ppm) | Phosphate<br>(0.4 ppm) | Ammonium<br>(0.6 ppm) | IBMWP<br>(93) | IMMi-T<br>(0.69) | IHF | QBR | AGR |
| --- | --- | --- | --- | --- | --- | --- | --- | --- | --- | --- |
| Ea | 8.01 ± 0.02 | 7.77 ± 0.27 | **30.55 ± 14.78** | 0.148 ± 0.102* | 0.050 ± 0.027 | 112 | 0.76 | 42 | 40 | 79.2 |
| Enc | 8.07 ± 0.08 | 9.20 ± 0.95 | 20.14 ± 9.61* | 0.009 ± 0.001 | 0.055 ± 0.032 | 134 | 0.91 | 67 | 50 | 46.6 |
| R1 | 7.92 ± 0.06 | 10.17 ± 0.35 | 0.95 ± 0.30 | 0.014 ± 0.003 | 0.187 ± 0.013 | 207 | 1.15 | 74 | 75 | 1.0 |
| R2 | 8.16 ± 0.08 | 8.80 ± 0.82 | 2.58 ± 0.78 | 0.010 ± 0.001 | 0.171 ± 0.015 | 129 | 0.93 | 71 | 50 | 17.6 |
| R3 | 8.38 ± 0.07 | 9.67 ± 0.68 | **41.85 ± 22.33** | 0.036 ± 0.016 | 0.069 ± 0.029 | 147 | 0.94 | 60 | 40 | 48.1 |
| Z1 | 8.11 ± 0.05 | 9.17 ± 0.33 | **88.83 ± 3.29** | 0.044 ± 0.035 | 0.135 ± 0.038 | **92** | **0.67** | 32 | 10 | 84.0 |
| Z2 | 8.22 ± 0.05 | 9.00 ± 0.15 | **84.68 ± 6.64** | 0.074 ± 0.040 | 0.019 ± 0.001 | 108 | 0.73 | 36 | 10 | 84.2 |
| T1 | 7.85 ± 0.13 | 6.93 ± 1.15* | 21.13 ± 12.35* | 0.136 ± 0.050 | 0.156 ± 0.031 | **59** | **0.35** | 29 | 10 | 70.0 |
| T2 | 8.16 ± 0.09 | 8.13 ± 1.21 | 19.77 ± 6.52* | 0.312 ± 0.141* | **1.584 ± 1.170** | **56** | **0.33** | 31 | 35 | 76.8 |
| Y1 | 8.41 ± 0.07 | 9.50 ± 0.40 | 0.63 ± 0.12 | 0.009 ± 0.001 | 0.165 ± 0.009 | 220 | 1.3 | 71 | 80 | 0.0 |
| Y2 | 7.52 ± 0.27 | 6.90 ± 0.64 | 4.12 ± 0.78 | 0.013 ± 0.003 | 0.159 ± 0.047 | 162 | 1.04 | 57 | 75 | 7.4 |

The average temperature (S3 Table) ranged from 9.7ºC at R1 and 15.5ºC at T2, while pH values were very similar across sites (ranging between 7.52 and 8.41). The average conductivity resulted lower than 500 µS/cm at Yalde (Y1 and Y2) and upper Reláchigo (R1 and R2). It considerably increased at all other sites, reaching values around 1000 µS/cm at lower Reláchígo (R3), Tuerto (T1 and T2), Zamaca (Z1 and Z2) and Ea streams and even higher values (i.e. mean conductivity 2077 µS/cm) at Encemero stream. Accordingly, exceptionally high values of calcium and sulfate were recorded at Encemero stream (sulfate = 896.79 ppm; calcium =575.52 ppm).

Among other studied ions, nitrate showed particularly high concentrations at different dates and sites. Considering average values, 4 sites exceeded 25 ppm, the highest value established by RD 817/2015 to consider a river in good ecological status (R3, Ea, Z1 and Z2; Table 1). However, there was a great variability in nitrate concentration among different dates (Table 1). Regarding the mean value of the rest of chemical variables included in the WFD, only ammonium exceeded the limits to reach good ecological status (site T2, Table 1), although phosphate concentrations exceeded the limits at T2 and Ea in May (0.68 and 0.44 ppm, respectively).

The IHF index differed among sites, with the lowest values found at Zamaca (Z1 and Z2), Tuerto (T1 and T2) and Ea streams (Table 1; S4 Table). The lowest values of these 5 sites respect to the others were recorded at components of IHF refers to substrate composition (component 3), velocity/depth regime (4) heterogeneity components (6), aquatic vegetation cover (7) and, especially, to embeddedness (1) (S4 Table). The value of this first component of IHF at Z1, Z2, T1, T2 and Ea was 0, since more than 60% of boulders, cobbles and pebbles were embedded in fine sediment. These sites, together with R3, also showed the lowest QBR values (Table 1; S4 Table). Macrophytes occupied between 30 and 60 % of the riverbed in every studied section, except at R2 (20%) and at Y1, Y2 and R1, where macrophyte cover was lower than 5%.

The IBMWP biotic index was above 93, the lowest value established by RD 817/2015 to consider a river at good ecological status, at every site except at Z1, T1 and T2 (Table 1). The highest IBMWP values were recorded in headwaters of Reláchigo and Yalde Streams (R1 and Y1), with values higher than 200. The IMMi-T was also below the good category threshold (0.70) at Z1, T1 and T2, and slightly higher values were recorded at Z2 and Ea. The highest values were once more determined at R1 and Y1 (Table 1).

Among land cover variables, the highest values of forest (coniferous and broadleaf) percentage upstream the studied sites were at Y1, Y2, R1 and R2 (i.e. higher than 30%; S5 Table). The area occupied by agricultural land upstream the river reach ranged from 0% at Y1 to 84.2 % at Z2. This variable was especially high (i.e. higher than 70%) at Ea, Z1, Z2, T1 and T2 (Table 1).

### Macroinvertebrate assemblage and agriculture pressures on streams

The two groups produced using cluster analysis (Fig 2) presented a 60 % of dissimilarity. Group 1 included the sites Enc, R1, R2, R3, Y1 and Y2, whereas Group 2 included the sites Ea, Z1, Z2, T1 and T2.

**Fig 2.**
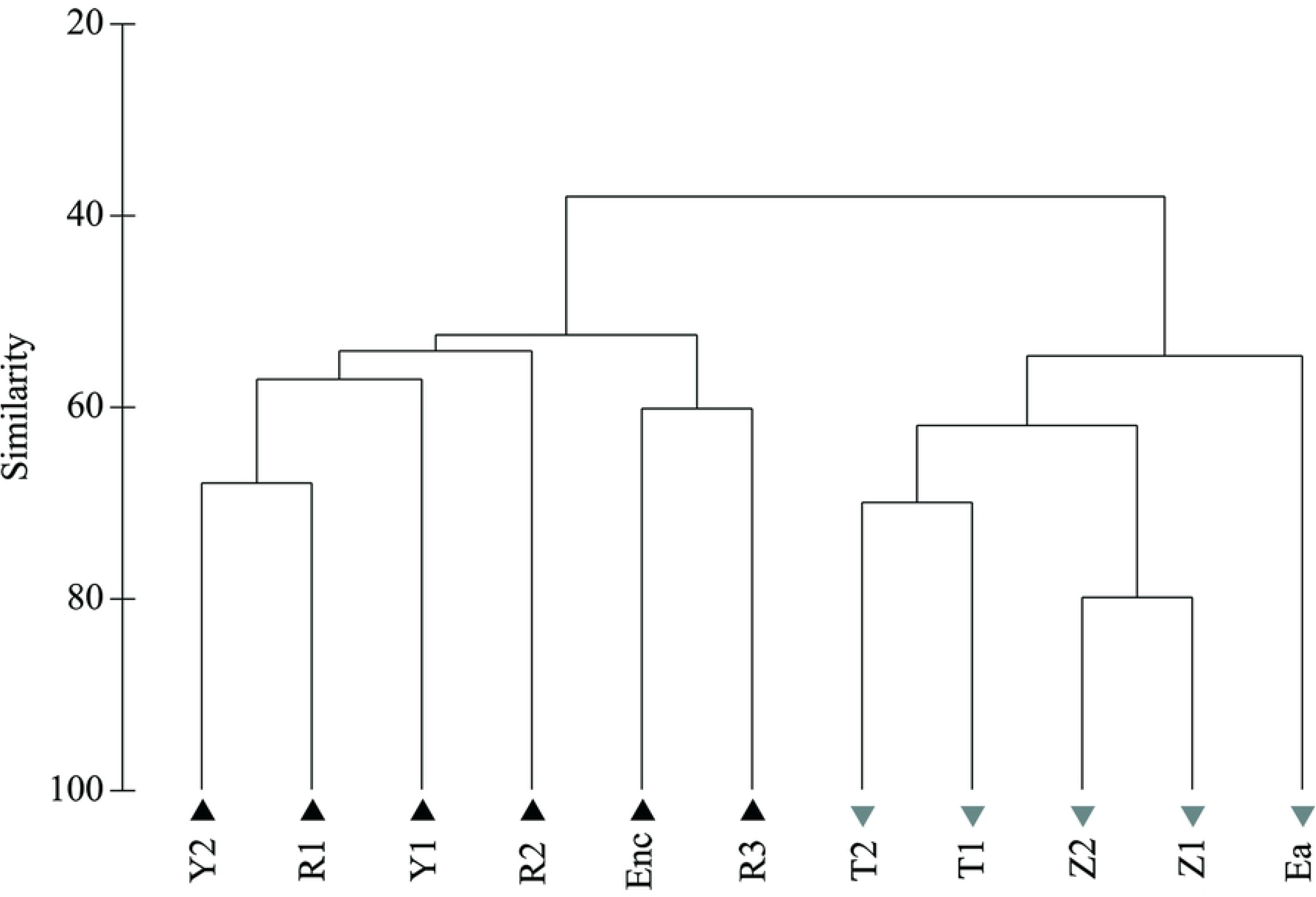
Hierarchical cluster. Hierarchical cluster analysis (UPGMA) based on the composition of the macroinvertebrate community of each site. The resultant groups are identified with black triangles (group 1) and grey inverted triangles (group 2).

The final model of the DISTLM explained 99.2% of the total variance and included the following significant explanatory variables (ranked in order of importance): total value of IHF index (43.9 % of total variance explained), percentage of area occupied by agricultural land upstream the river reach (11.8 %), nitrate concentration (9.4 %), the ratio of valley width to channel width (8.6 %) and the component of the QBR index referring to channel alterations (7.3 %). Other variables included in the final model were not significant, although they also contributed to explain the model variance. They were the component of the QBR index refers to riparian vegetation cover (7.2 %), percentage of land occupied by pasture upstream (4.3 %), chloride concentration (4.2 %) and macrophyte cover (2.2 %) (Fig 3).

**Fig 3.**
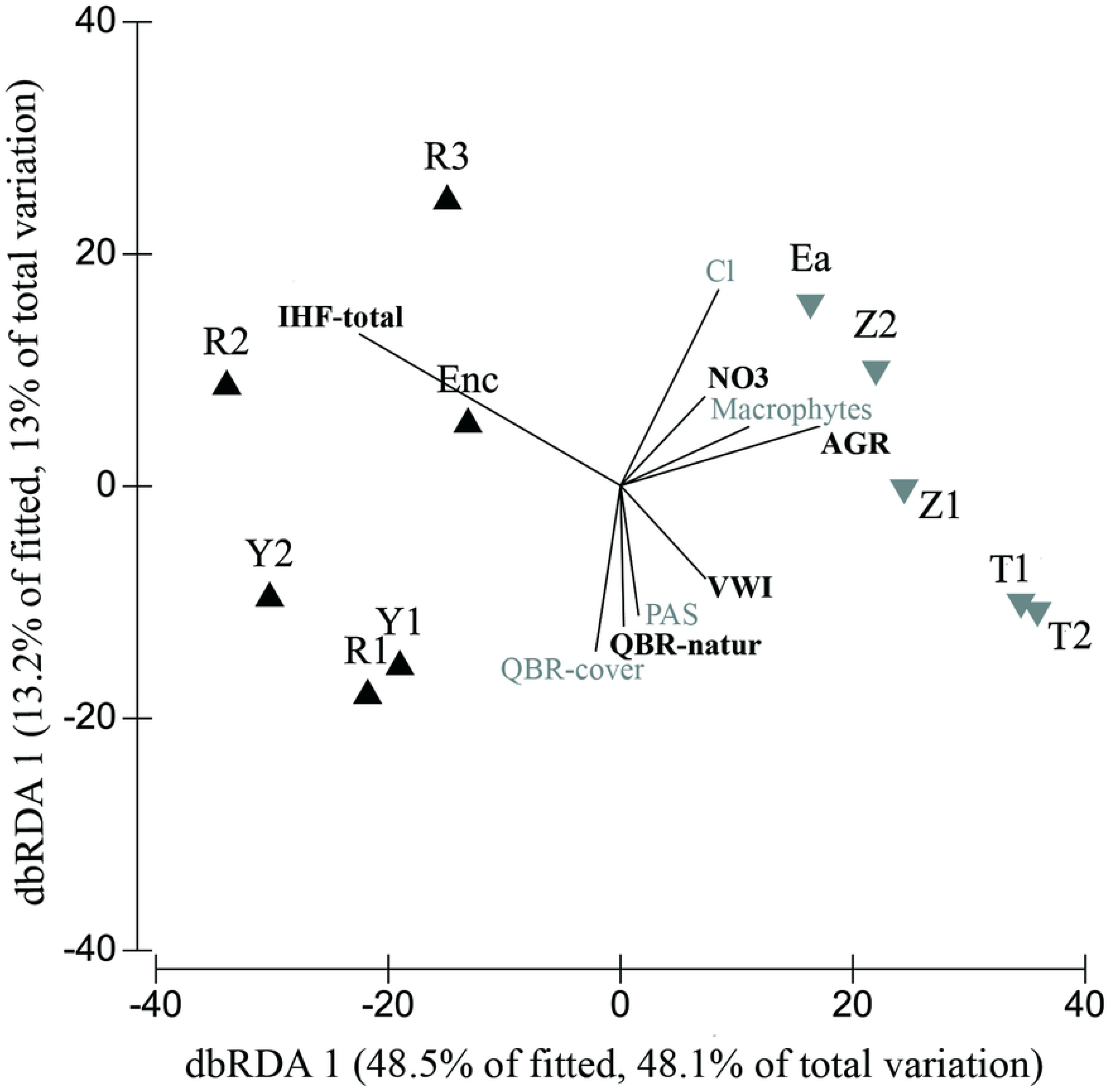
DISTLM analysis. Step-wise DISTLM analysis using AIC as selection criterion for the selection of explanatory variables. Only those variables included in the final model are plotted, and those significantly explaining the macroinvertebrate community composition (P<0.05) are highlighted in bold. Sampling sites previously grouped by cluster analysis are shown with different symbols (black triangles: group 1; grey inverted triangles: group 2). IHF: Total score of the Habitat Fluvial Index; QBR-natur: component of QBR referred to river channel alteration; NO3: nitrate concentration; Cl: chloride concentration AGR: area occupied by agricultural land upstream the river reach; VWI: Valley Width Index or ratio of valley width to channel width; PAS: area occupied by pasture upstream the river reach.

Group 1 was mainly characterized by EPT taxa such as Polycentropodidae, Heptageniidae, Leptophlebiidae or Leuctridae, which are sensitive to pollution according to the IBMWP (Table 2). Other taxa associated to this group were Gammaridae, Ceratopogonidae or the genus *Esolus* and *Elmis* (Elmidae). The group 2, affected by a greater agricultural surface, was associated with taxa with lower IBMWP score (Table 2), like Oligochaeta and Chironomidae, together with other dipterans like Simuliidae or Psychodidae and other taxa such as Hydroptilidae or Haliplidae.

**Table 2.**
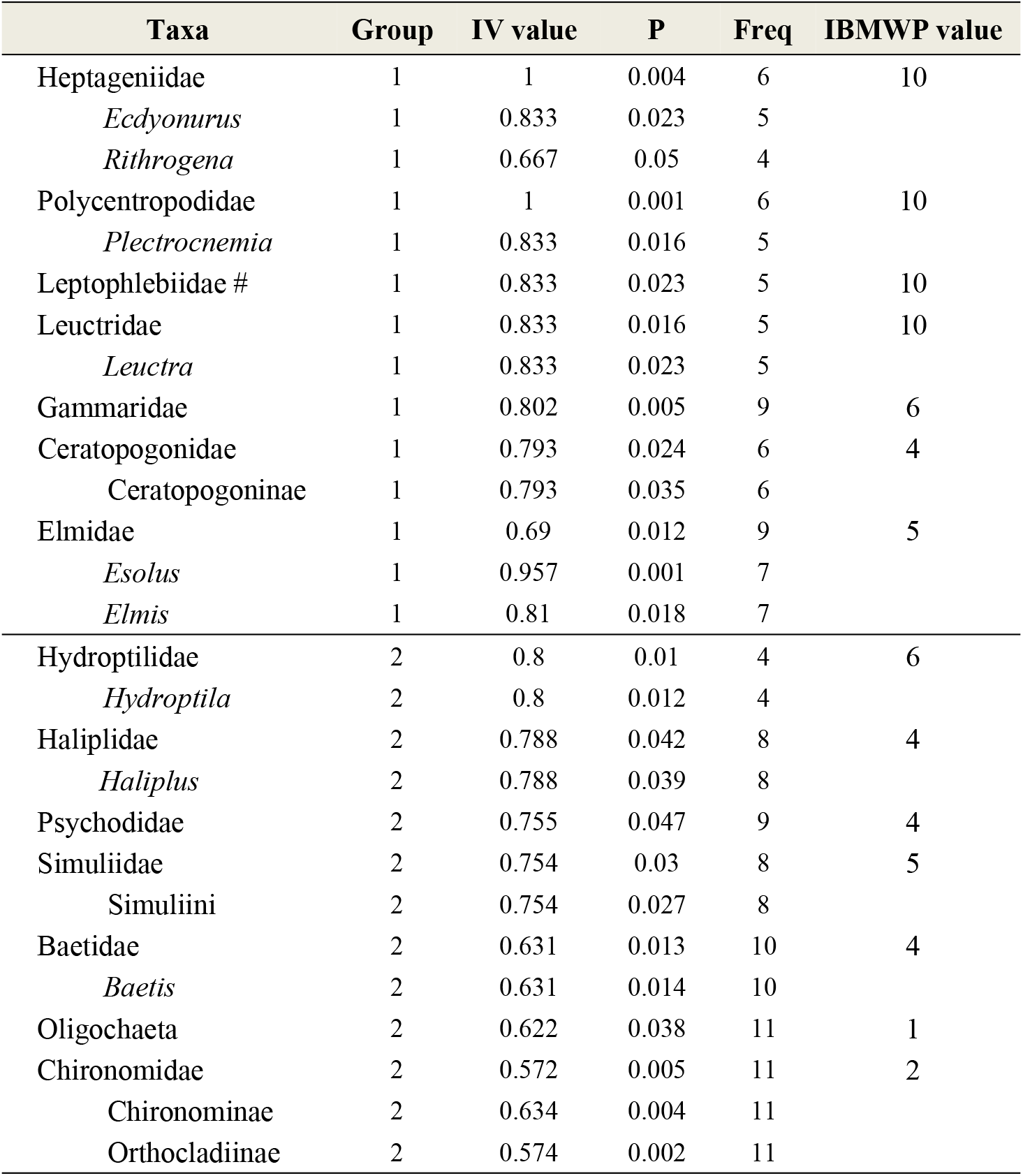
IndVal analysis. IndVal analysis results showing taxa significantly associated with one of the previous established groups. For those taxa included in the IBMWP index, their indicator value is shown. # means that the whole family is significantly different between groups, but no differences were found at the genus level.

Biological trait categories significantly differed between groups 1 and 2 (Fig 4). Group 1 was characterized by crawlers living on large substrate, shredder and scrapers feeding microphytes, taxa adapted to oligotrophic conditions, gill and plastron respiration and brief reproductive cycle. Group 2 was characterized by taxa living and feeding on fine sediment and macrophytes, adapted to eutrophic conditions and with longer reproductive cycle (Fig 4).

**Fig 4.**
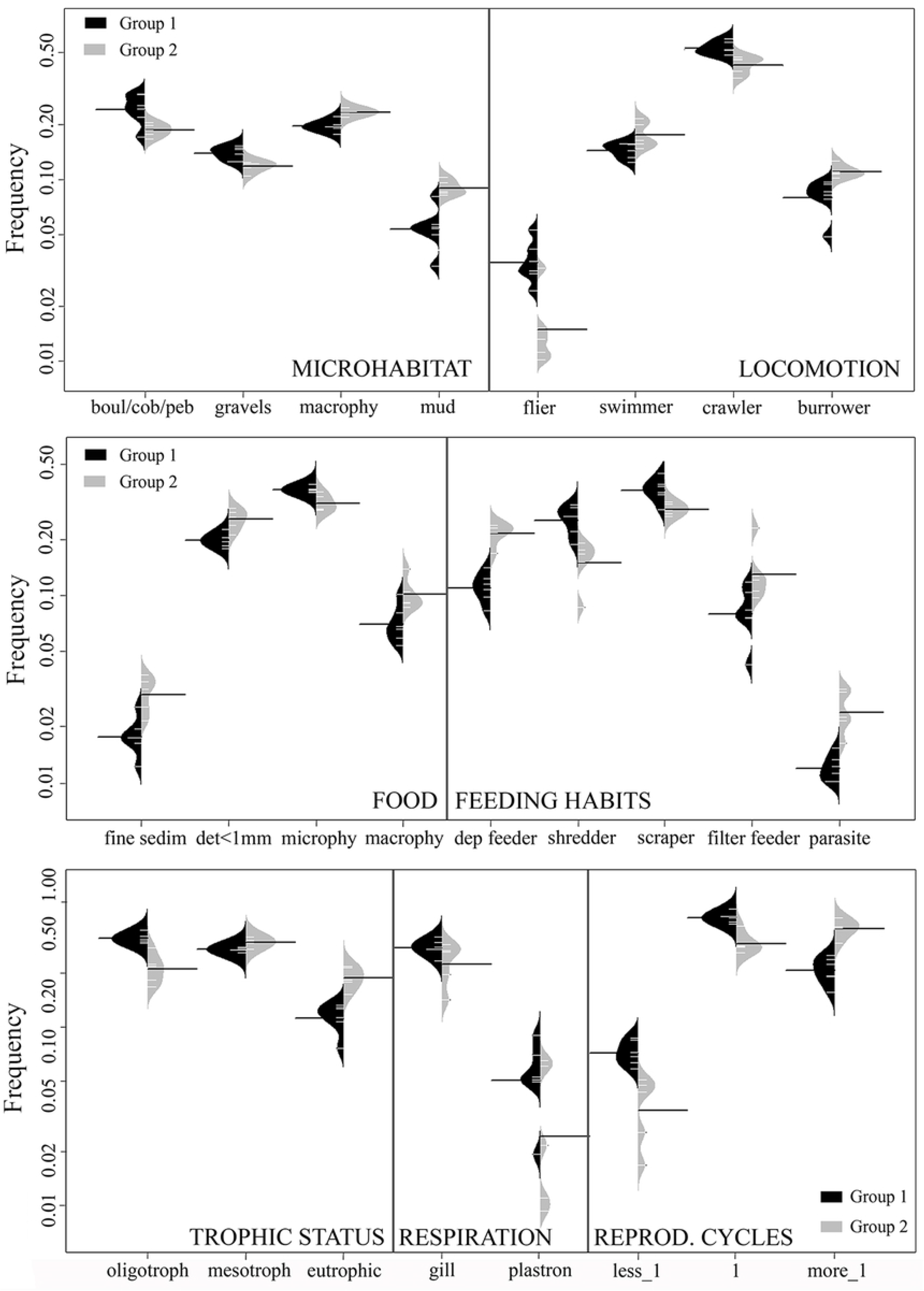
Biological traits analysis. Asymmetric beanplots of the biological trait categories significantly different (Mann Whitney U test, P<0.05) between previously established groups. The individual observations of each site are shown as small white lines and the average of each group is shown as black line.

## Discussion

Agricultural activity clearly altered the stream biota in the study area. In agreement with previous studies [13,15,17,24] agricultural intensification led to a replacement of EPT taxa by tolerant taxa such as Oligochaeta or Chironomidae. Sites with higher upstream area occupied by agricultural land were dominated by taxa with short generation times, a biological trait that has been associated with resilient strategies in human impacted sites [27].

Nitrate significantly contributed to alter macroinvertebrate assemblages in the studied streams, as it has been observed in other agricultural areas [15,21]. Intensive agriculture reduced the percentage of oligotrophic macroinvertebrates. Nitrate is able to convert oxygen carrying pigments of aquatic animals to forms that are incapable of carrying oxygen (e.g. methemoglobin; [19,48]). In this regard, the mean nitrate concentration was enough to adversely affect, at least during long-term exposures, freshwater invertebrates (i.e. 10 mg/L, according to Camargo *et al.* [19] and Pearson *et al.* [21]) at 7 of the study sites. However, nitrate uptake in aquatic animals is limited [49] and nitrate impact on ecosystems can be related to other processes, like an increase in autotrophic production [11]. Accordingly, macrophyte cover increased in the most agricultural impacted sites, associated with high nitrate content and reduced shading due to riparian forest degradation. Thereby, agricultural impact favored taxa living and feeding on macrophytes and those adapted to eutrophic conditions.

The variation in aquatic macroinvertebrate assemblage among sites differentially affected by agriculture in the present study was best explained by variables associated with hydromorphological alteration of the stream habitat and topography. Specifically, the total IHF index, the QBR components that refer to riparian vegetation cover and river channel alteration and the ratio of valley width to channel width (VWI) are responsible of 67% of macroinvertebrate assemblage variation among sites. Accordingly, different authors have pointed out the physical habitat degradation as the major threat in many agricultural watersheds and not water pollution. For instance, Genito *et al.* [12] suggested that a high percentage of agricultural land cover led to a macroinvertebrate assemblage composition that reflected stream habitat degradation, while nitrate concentration had a weaker association with assemblage alteration. Shields *et al.* [50] pointed out the importance of stream habitat conservation in agricultural areas, especially related to channel degradation, large wood removal or increase of fine sediments deposition. Accordingly, fine sediment has been determined as a more pervasive stressor to macroinvertebrate assemblage than augmented nutrient concentrations in streams around agricultural areas [25] and in stream mesocosm experiments [17]. Beyond agricultural activity, several studies have confirmed the strong impacts of fine sediment on stream invertebrates (e.g. [51–54]).

Intensive agriculture in the study area alters the riverbed, simplifies the river habitat conditions -e.g. high substrate embeddedness in fine sediment or low velocity regimes and elements heterogeneity- and profoundly deteriorates the riparian forest (Fig 5). According to Genito *et al.* [12] and Piscart *et al.* [55], a reduction of riparian forests can decrease the abundance of shredders leading to a lower litter breakdown rate. Riparian alteration also enhances, together with the intensive soil tillage, the amount of sediment entering the streams [11], which represented one of the most important variables altering the functional structure of macroinvertebrate assemblages in our study sites. Increased stream sedimentation has often been associated with nonpoint sources arising from agricultural land uses and is mainly related to the loss of riparian vegetation through direct (inputs) and indirect (flow-mediated) effects on deposited sediment [25].

**Figure 5.**
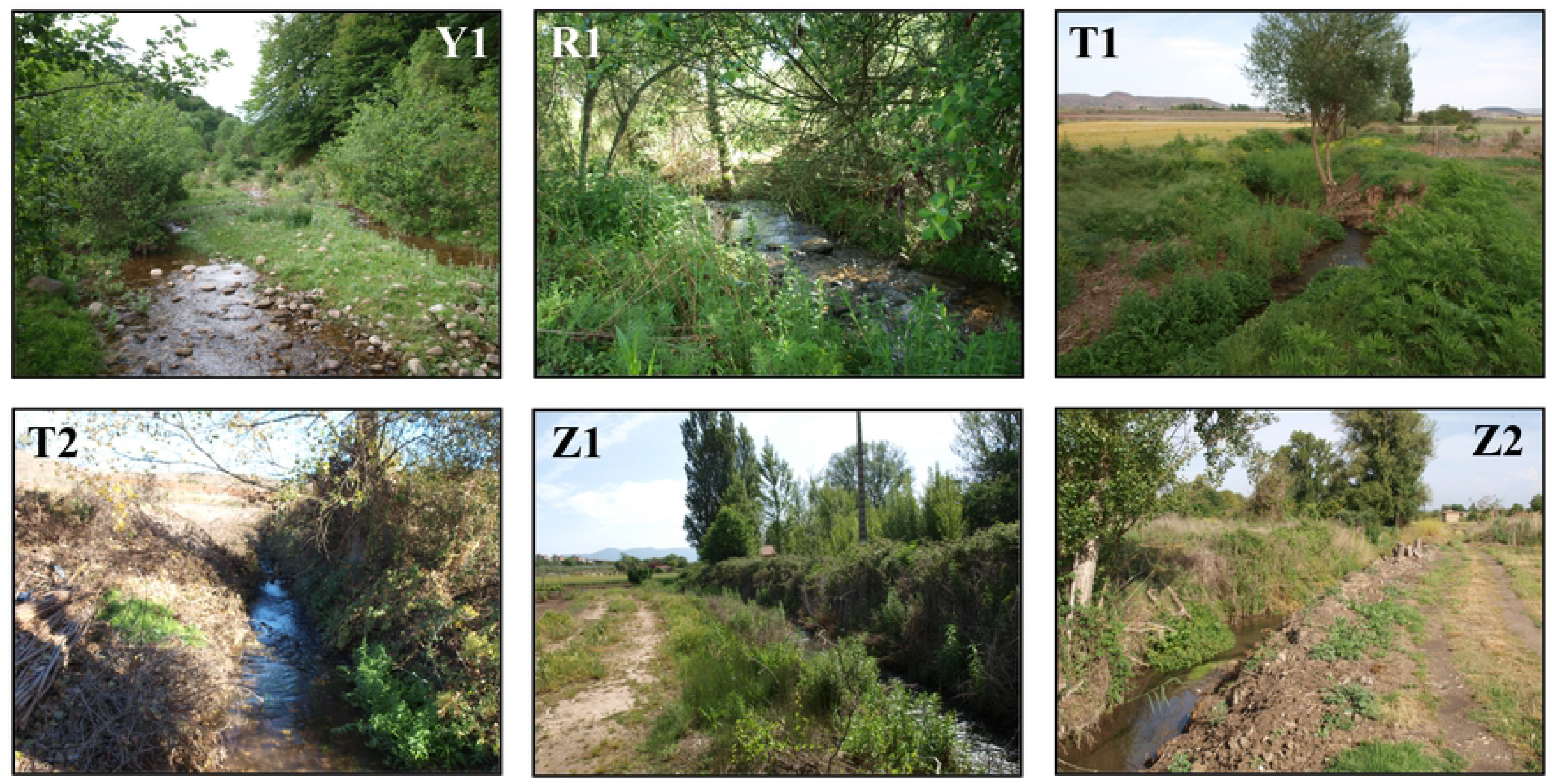
Photographs of the studied sites with the highest (white acronyms) and the lowest (black acronyms) QBR and IHF total values. Y1 (QBR=80; IHF=71), R1 (75; 71), T1 (10; 29), T2 (35; 31) Z1 (10; 32), Z2 (10; 36).

In the present study, agricultural activity was associated with a higher proportion of boulders, cobbles and pebbles embeddedness in fine sediment (component 1 of the IHF index). This could explain the reduction in organisms adapted to move on large substrate (i.e. crawlers), since fine sediment decreases habitat availability [25]. On the contrary, the percentage of invertebrates living in the mud and burrowers increased, as it has been previously shown for sediment input increases in rivers [17,28]. Agricultural intensification also favored organisms that fed on fine sediment, while negatively affected scrapers, since the sediment covers larger substrate reducing microphytes availability for stream fauna. Finally, taxa breathing by plastron and gills were significantly less frequent in sites with higher agricultural surface. For example, we observed a decrease of *Elmis* and *Esolus*, which have larvae with anal gills and aquatic adults with plastron respiration [41,56]. Concordantly, species of Coleoptera relying on a bubble or plastron for breathing have been shown to be sensitive to sediment increases [57].

As a consequence of the intense stream alteration associated with agriculture in the study area, several sites did not meet the legal requirements to be considered in good ecological status according to the WFD. Three sites did not reach the good ecological status based on IBMWP and IMMi-T biotic indices and nitrate exceeded the “good” limits at 7 sites, despite all of them, except Z1, were outside of NVZs. However, the WFD indicators seem to underestimate hydromorphological alterations, since they are not specifically considered to determine the good ecological status of the water bodies. According to Houlden [58], hydromorphology is a “supporting element” in the WFD, which is only used to confer the high ecological status. This has caused that, despite the key role of hydromorphological alteration caused by agriculture in the study area, habitat altertion has been largely overlooked by water managers within the Ebro basin [59]. The official reports point to nitrate as the main agricultural pressure to the studied streams but they only suggest the implementation of “good agricultural practices” at a few sites, and these practices are not detailed [59].

## Conclusions and recommendations

Aquatic macroinvertebrates are a good indicator of agricultural impacts, since both the taxonomic composition and the functional structure responded to agricultural intensification. In agreement with Magbanua *et al.* [15], the study of the functional structure was key for understanding the consequences of agricultural practices on stream communities. Intensive agriculture around NVZs in the study area significantly altered the stream habitats and its associated macroinvertebrate assemblages. High nitrate content is partially responsible for such alteration, but hydromorphological impacts, especially the increase in the amount of sediment that enters the streams, seem to be the main driver behind stream degradation.

The implementation of the ND and the WFD are not enough to prevent the degradation of streams in agricultural areas. We suggest, as other authors did [30,60], to extend the NVZs in the study area, mainly “Zamaca and Oja alluvial”, since nitrate exceeded the legal limits in 7 studied sites to achieve good ecological status (despite 6 of them are located upstream of NVZs). Moreover, farming practices established by regional administration in NVZs have not been effective to reduce nitrate pollution even in these areas [61], so the compliance of these practices should be more rigorously controlled. At the same time, water quality monitoring programs are not able to identify the hydromorphological pressures existing in the area. A proper monitoring of these pressures should be designed, and adequate practices should be implemented to achieve a good ecological status. We recommend dredging and channelization restrictions, protection of riparian vegetation, and “Good agricultural environmental conditions” (e.g. reduced tillage, winter cover crops, plant residues, stone walls or grass margins and contouring; [62]), which has reduced soil loss from European arable lands by around 20% in the past decade [63].

Finally, it should be noted that the recovery of stream ecosystems in agricultural areas requires not only mitigation actions but also restoration to accelerate ecological recovery [64]. It is necessary to carry out restoration plans, at least in Ea, Zamaca and Tuerto Streams, which were the most hydromorphologically impacted. Among the restoration strategies available to improve ecological status, the establishment of riparian buffers has been the most frequently and successfully used to improve stream habitat [64–68], which should try to include a minimum perimeter of 20-30 m along the impacted sites [64].

## Supporting information

**S1 Table. Environmental variables studied.** Environmental variables studied in each site, categories and codes used in the present study.

**S2 Table. Biological traits studied.** Biological traits and categories studied according to Tachet *et al*., (2006)

**S3 Table. Results of physicochemical variables.** Average values (four sampling dates) ± standard error (SE) of physicochemical variables not included in Table 1.

**S4 Table. Results of hydromorphological variables.** Values of hydromorphological variables not included in Table 1.

**S5 Table. Results of topographic, geologic and land cover variables.** Values of topographic, geologic and land cover variables not included in Table 1. Note that the percentages do not have to sum 100%

